# Distinct blood DNA methylation profiles in subtypes of orofacial cleft

**DOI:** 10.1101/113266

**Authors:** Gemma C Sharp, Karen Ho, Amy Davies, Evie Stergiakouli, Kerry Humphries, Wendy McArdie, Jonathan Sandy, George Davey Smith, Sarah Lewis, Caroline L Relton

## Abstract

**Background:** There is evidence that different subtypes of orofacial cleft have distinct aetiologies, although the precise molecular mechanisms underlying these are unknown. Given the key role of epigenetic processes such as DNA methylation in embryonic development, it is likely that aberrant DNA methylation may also play a part in the development of orofacial clefts.

**Methods:** In this study, we explored whether blood samples from children with different cleft subtypes showed distinct DNA methylation profiles.

In whole blood samples from 150 children from the Cleft Collective cohort study, we measured DNA methylation at over 450,000 sites on the genome. We then carried out epigenome-wide association studies (EWAS) to test the association between methylation at each site and cleft subtype (cleft lip only CLO n=50; cleft palate only CPO n=50; cleft lip and palate CLP n=50).

**Results:** We found four genomic regions differentially methylated in CLO compared to CLP, 17 in CPO compared to CLP and 294 in CPO compared to CLO. These regions included several mapping to genes that have previously been implicated in the development of orofacial clefts (for example, *TBX1, COL11A2, HOXA2, PDGFRA*) and over 250 novel associations.

**Conclusion:** Our finding of distinct methylation profiles in different cleft subtypes might reflect differences in their aetiologies, with DNA methylation either playing a causal role in development of OFC subtypes or reflecting causal genetic or environmental factors.

## Background

Orofacial clefts (OFCs) are a set of common birth defects that affect roughly 15 in every 10,000 births in Europe [1]. A child born with an OFC may face difficulties with feeding, speech, dental development, hearing and social adjustment. At considerable health, emotional and financial costs, they undergo surgery in the first year of life and many need additional surgical procedures later in life. They may experience low self-esteem, psychosocial problems and poor educational attainment, and the condition can harm the emotional wellbeing of the whole family [2–4],

Syndromic OFCs are often known to be caused by a specific genetic or chromosomal anomaly, whereas non-syndromic cases, which comprise around 70% of cases of cleft lip with or without cleft palate, have a complex aetiology involving both genetic and environmental factors [2], Furthermore, there is increasing evidence that the three main subtypes of OFC, cleft palate only (CPO), cleft lip only (CLO) and cleft lip with cleft palate (CLP) (Figure 1), are aetiologically distinct. For example, there is a higher risk of familial recurrence of the same subtype compared with risk of recurrence of a different subtype [5]. CLO tends to be combined with CLP in research because it has traditionally been thought of as a less extreme manifestation of CLP. However, differences in sex ratios and rates of twinning and consanguinity between the two subtypes provide some evidence that CLO and CLP may be aetiologically distinct [6].

**Figure 1.**
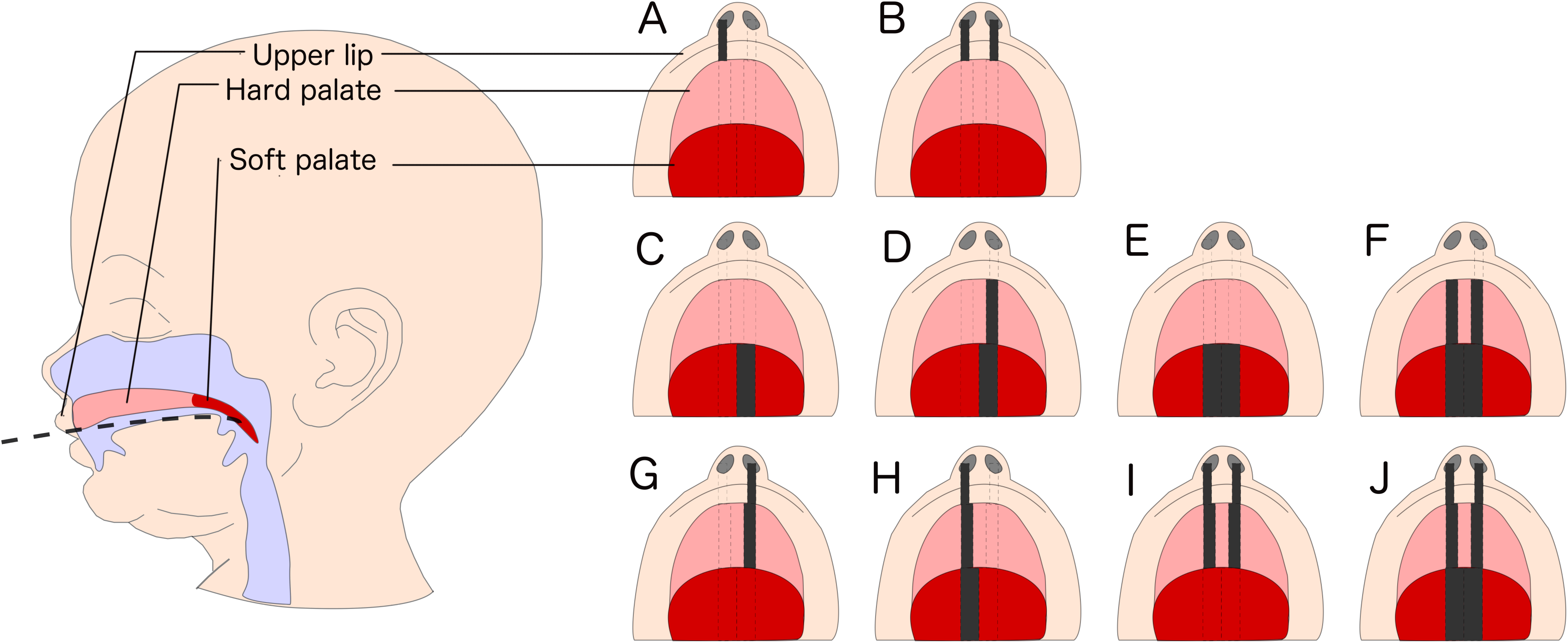
Orofacial cleft subtypes. Orofacial clefts are traditionally categorised as either cleft lip only (CLO; A, B), cleft palate only (CPO; C-F) or cleft lip with cleft palate (CLP; G-J). Further subtyping can be made according to laterality and whether the soft and/or hard palate is affected.

If CPO, CLO and CLP have distinct aetiologies, the precise underlying molecular mechanisms are currently unknown. Given the key role of epigenetic processes such as DNA methylation in embryonic development, we and others have hypothesised that aberrant epigenetic processes may also play a role in the development of OFCs [2,7,8]. This hypothesis has been supported by data suggesting an important role for DNA methylation and other epigenetic processes in regulating normal orofacial development and OFCs in mice [9–14], but published epigenetic data for OFCs in humans is lacking.

A clearer understanding of the aetiologies of OFCs will help to inform better strategies for screening, diagnosis, counselling and prevention. Therefore, we carried out this study to explore distinct aetiologies of OFC subtypes, by generating and examining DNA methylation profiles in whole blood samples from non-syndromic children with CLO, CPO and CLP.

## Methods

### Participants

Participants were children from the United Kingdom enrolled in the Cleft Collective birth cohort study between 2013 and 2016[15–16]. Families of a child with an OFC were invited to take part soon after the child was born. Demographic and lifestyle information for both parents was collected via questionnaire. Blood samples were collected at time of surgery to repair the OFC. Additional details on the surgery and OFC were collected on a surgical form. For the purposes of the current study, a sample of 150 believed-to-be non-syndromic children were randomly selected. The sample was stratified by OFC subtype (CLO, CPO, CLP) resulting in 50 children per group.

### Classification of OFC

Details on the cleft phenotype were collected from surgical forms completed at the time of operation and from parental questionnaires. Surgeons recorded the phenotype using either the LAHSAL or LAHSHAL classification [17], which was condensed to CPO, CLP or CLO for the purposes of this study. Parents used this simplified classification of subtype (CPO, CLP or CLO). Where data were available from both sources, we compared the reported subtype and found no discrepancies.

### Other variables

The child′s age at biological sample collection was calculated from the child′s date of birth and date of surgery. We also predicted child′s ′epigenetic age′ using the method developed by Horvath [18] (discussed in more detail in Additional File 1). ′Age acceleration′ was calculated as the residuals from a linear regression of epigenetic age on actual age at sample collection. A positive value corresponds to an individual whose epigenetic age is ahead of their actual age, and vice-versa. Sex was initially assumed by staff at the Cleft Collective using the child′s name and later confirmed by parental questionnaires, where available, and NHS Digital data if explicit consent was held. Mothers self-reported how much they smoked around the time of conception, and this was classified for the purposes of this study as any or no smoking around conception. Additionally, a score to predict in utero/early-life smoke exposure was calculated from the child′s blood DNA methylation data. The score was calculated as previously described [19] using a weighted sum of methylation beta values at 26 maternal-smoking-associated methylation sites identified in cord blood [20]. The efficacy of this score for predicting maternal smoking is discussed in Additional File 1. Information on the mother′s occupation was dichotomised as either non-manual skilled work, or manual/unskilled/no work. Information on mother′s education was dichotomised as achieving a university degree/above, or not achieving a university degree. Information on parity was dichotomised for this study as no previous children or one or more previous children. Maternal and paternal age in years were reported by the mother and treated as continuous variables. Maternal and paternal ethnicity were reported by the mother and used to deduct child ethnicity as white or other. For each model, surrogate variables were generated using the sva package [21,22] in R [23] to capture residual variation associated with technical batch and cellular heterogeneity. The number of surrogate variables (10) was estimated by the sva algorithm using the methylation data and the model matrices. Blood cell type proportions were also estimated using the Houseman method [24,25] for use in a sensitivity analysis.

### DNA methylation

Upon arrival at the Bristol Bioresource Laboratories (BBL), whole blood samples were immediately separated by centrifugation into white blood cell and plasma aliquots before storage at −80°C. DNA from the white blood cells was extracted and genome-wide DNA methylation was measured using the Illumina Infinium HumanMethylation450 BeadChip platform. Data were pre-processed in R version 3.3.2 with the meffil package [26], Functional normalisation [27] was performed in an attempt to reduce the non-biological differences between probes. Of the original 150 samples, three failed quality control due to a mismatch between reported and methylation-predicted sex (and additional data from NHS Digital or parental questionnaire was not available to cross check). In addition, we removed 944 probes that failed quality control in meffil and a further 1,058 probes that had a detection P-value >0.05 for >5% of samples. Finally, we removed 11,648 probes mapping to the X or Y chromosomes and 65 SNP probes included on the array for quality control purposes. This left 472,792 probes in the dataset for further analysis. Extreme outliers in the methylation data were identified using the Tukey method (<l^st^ quartile-3*IQR; >3^rd^ quartile+3*IQR) and set as missing. The median number of samples removed per probe was 0 (IQR: 0 to 1; range 0 to 72).

### Statistical analysis

We assessed the association between parental and child characteristics and cleft subtype using chi-squared or t-tests. We also used linear regression to explore whether 'epigenetic age' (age predicted using the methylation data) differed from true age at sampling and whether any deviation (age acceleration) was associated with OFC subtype or any parental characteristics.

Epigenome-wide association studies (EWAS) were conducted in R version 3.3.2 [23]. For our main EWAS analyses, we used linear regression to model cleft subtype as the exposure and untransformed methylation beta values as the outcome. To identify blood methylation profiles specific to each subtype, we made three pairwise comparisons: CPO compared to CLP (CPOvsCLP), CLO compared to CLP (CLOvsCLP), CPO compared to CLO (CPOvsCLO). All models were adjusted for sex because previous studies have found different sex ratios for OFC subtypes. In order to adjust for technical batch effects and cellular heterogeneity, we calculated surrogate variables and included these in all models. We also conducted a sensitivity analysis using chip ID to adjust for batch and Houseman-estimated cell proportions to adjust for cellular heterogeneity. For results from the CPOvsCLO and the CPOvsCLP EWAS analyses we removed age-related CpGs as described below. P-values were corrected for multiple testing using the Bonferroni method and a threshold of 0.05, i.e. an uncorrected P-value threshold of 1*10^−7^. Regression coefficients are interpreted as the difference in mean methylation beta value in children with one subtype compared to children with another subtype.

In addition to the EWAS analyses at individual CpGs, we also used Comb-P [28] to detect differential methylation across larger regions of the genome. This approach is statistically more powerful and has been associated with a lower rate of false positive findings compared to EWAS at individual CpGs [29] Using genomic location and P-values from our individual CpG EWAS results, Comb-P identifies regions that are enriched for low P-values. It then calculates and adjusts for auto-correlation between those P-values using the Stouffer-Liptak-Kechris correction and performs Sidak correction for multiple-testing. Differentially methylated regions (DMRs) were defined as regions fulfilling these criteria: 1) contains at least two probes, 2) all probes within the region are within 1000 base pairs of at least one other probe in the region, 3) the Sidak-corrected P-value for the region is <0.05.

### Special consideration of age at sampling

Blood samples were collected at first surgery, which is typically around 3-6 months after birth for lip repair and 6-18 months after birth for palate repair. Therefore, we anticipated that the children with CLO and CLP would be younger than the children with CPO. Previous studies have shown that age, particularly during this early developmental period, is strongly associated with methylation [30–32]. We refrained from adjusting for age at sampling because it is not a true confounder (it cannot plausibly cause OFC subtype). Instead, we considered it a nuisance variable and dealt with it by ‘filtering out’ any age at sampling-related CpGs from our main analysis. To do this, using all the participants in our sample, we ran an EWAS of age at blood sampling, and for any age-associated CpGs (uncorrected P-value <0.05), we set the EWAS P-values from the CPOvsCLP and the CPOvsCLO analyses to 1. This meant that we were filtering out age-related CpGs from our main EWAS results while maintaining the same multiple testing burden and array structure for the region-based analysis. In the age at sampling EWAS, child's age in months was modelled as the exposure with methylation as the outcome. The model was adjusted for 10 surrogate variables for technical batch and cellular heterogeneity. We confirmed that our age-at-sampling EWAS was effectively identifying age-related CpGs (independent of OFC subtype) by inspecting heterogeneity statistics from a meta-analysis of three separate age-at-sampling EWASs run within each OFC group (more details in Additional File 1).

### Functional analysis

To explore the function of any OFC-associated DMRs, we used the missMethyl[33] R package to test for enrichment of any gene ontology (GO) classification terms or Kyoto Encyclopaedia of Genes and Genomes (KEGG) pathways. This method corrects for biases in the genomic coverage of the Illumina Infinium HumanMethylation450 BeadChip array. We also looked up gene annotations from our DMRs in recently curated lists of OFC-related genes from 1) the DisGeNET database of diseases and related genes from human, rat and mouse studies [34] and 2) a bioinformatics study of OFC-related genes in human and animal studies published by Funato et al. [35].

## Results

### Characteristics of participants

From 150 participants selected for the study, 147 passed quality control for methylation data and 146 had information on all variables in the main model (subtype, age at sampling, sex). Participant characteristics are summarised in Table 1. As we expected, children with CPO were on average seven months older than children with CLO (t-test P=1.3*10^−19^) and six months older than children with CLP (t-test P=35.8*10^−14^) because of the timing of surgery for lip and palate repair (Figure 2). Accordingly, the same pattern was seen for epigenetic age, despite weak correlation with actual age (Spearman's rho for all participants 0.7; CPO 0.5, CLO 0.3, CLP 0.3). Participants with CLP tended to have a higher epigenetic age than their actual age (mean residual 0.8 months) whereas participants with CLO or CPO tended to have a lower epigenetic age than their actual age (mean residual −0.3 and −0.5, respectively). However, the confidence intervals crossed the null and t-test P-values for differences between subtypes were large (ranging 0.1 to 0.9). Age acceleration was also not associated with any measured confounder (Additional File 1). Participants with CLP were more likely to be male than participants with CLO (chi-squared P=0.02) or CPO (chi-squared P=0.002), but there was no difference in the sex ratio between participants with CLO and CPO (chi-squared P=0.60). According to maternal self-report of smoking behaviour, participants with CPO were more likely to have mothers who smoked around the time of conception compared to participants with CLP and CLO (chi-squared p-value=0.07). It should be noted that there was a particularly high level of missing data for this variable (70%). A tobacco exposure score calculated from the blood DNA methylation data was not associated with cleft subtype (chi-squared P=0.361).

**Figure 2.**
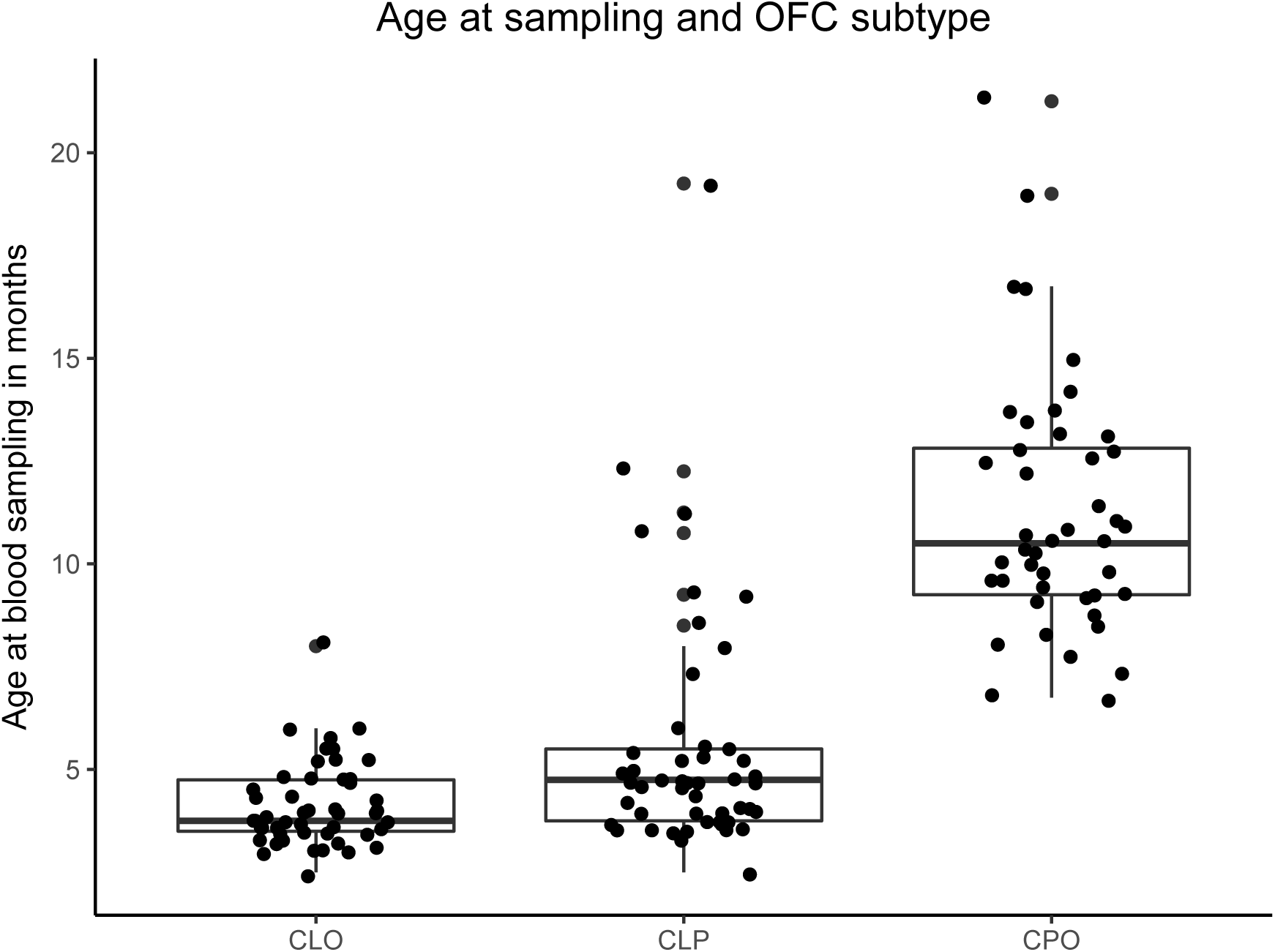
Age at sampling and OFC subtype. Children with CPO were older on average than children with CLO or CLP because surgery for palate repair usually occurs later than surgery for lip repair.

**Table 1.** Participant characteristics.

|  | **CLO (n=49)** | **CLP (n=49)** | **CPO (n=49)** | **P-value<sup>a</sup>** |
| --- | --- | --- | --- | --- |
| **Age in months at sample collection (95% CI)** | 4.1 (3.8, 4.4) | 5.5 (4.7, 6.3) | 11.2 (10.3, 12.0) | $3.9 \times 10^{-27}$ |
| **Epigenetic age<sup>b</sup> in months at sample collection (95% CI)** | 5.4 (4.2, 6.5) | 8.7 (7.1, 10.3) | 13.7 (11.9, 15.5) | $5.0 \times 10^{-11}$ |
| **Age acceleration<sup>c</sup> in months (95% CI)** | -0.3 (-1.5, 0.8) | 0.8 (-0.1, 1.8) | -0.5 (-2.0, 1.0) | 0.23 |
| **Female (%)** | 19 (39%)<br>(1 missing) | 8 (16%) | 23 (47%) | 0.004 |
| **White ethnicity (%)** | 25 (93%)<br>(22 missing) | 27 (93%)<br>(20 missing) | 23 (88%)<br>(23 missing) | 0.80 |
| **Maternal age at conception (95% CI)** | 30.9 (29.4, 32.4) | 29.4 (27.9, 30.9) | 31.1 (29.8, 32.4) | 0.21 |
| **Paternal age at conception (95% CI)** | 34.1 (32.2, 35.9) | 32.7 (30.8, 34.7) | 35.3 (33.4, 37.1) | 0.23 |
| **Methylation-predicted tobacco exposure score (95% CI)** | 0.004 (-0.1, 0.1) | -0.1 (-0.2, 0.1) | 0.1 (-0.1, 0.2) | 0.361 |
| **Self-reported maternal smoking around** | 0 (0%)<br>(33 missing) | 4 (36%)<br>(38 missing) | 7 (44%)<br>(33 missing) | 0.01 |
| **conception (%)** |  |  |  |  |
| **Maternal education:**<br>**university degree or**<br>**higher (%)** | 13 (48%)<br>(22 missing) | 13 (48%)<br>(22 missing) | 15 (58%)<br>(23 missing) | 0.73 |
| **Maternal occupation:**<br>**non-manual work** | 10 (38%)<br>(23 missing) | 13 (50%)<br>(23 missing) | 14 (61%)<br>(26 missing) | 0.29 |
| **Parity >=1** | 12 (43%)<br>(21 missing) | 19 (66%)<br>(20 missing) | 11 (46%)<br>(25 missing) | 0.18 |

<sup>a</sup>P-values were calculated using either ANOVA or chi-squared/Fishers tests
<sup>b</sup>Epigenetic age is age predicted using DNA methylation as described in Horvath et al. [18]
<sup>c</sup>Age acceleration refers to the residuals from a linear regression of epigenetic age on actual age as described in Horvath et al. [18]

### Individual CpG epigenome-wide study

After Bonferroni correction for multiple testing, there were no CpGs where DNA methylation in blood was associated with either CLO (n=48) compared to CLP (n=49), or CPO (n=49) compared to CLP (P>1*10^− 7^). In contrast, 335 CpGs were associated with CPO compared to CLO (Additional File 2, Table S1). Sensitivity analyses adjusting for chip ID and Houseman-estimated cell types (instead of surrogate variables) did not yield substantially different results (Additional File 1). We considered that some of these associations might be better explained by differences in age than OFC subtype, so we compared results to those of our EWAS of age at sampling (described above). There were 29,984 CpGs associated with age at sampling with an uncorrected P-value<0.05 (N participants in analysis 139; Additional File 2 Table S2). Confidence that these CpGs are truly associated with age (independently of OFC subtype) comes from our observation of low heterogeneity when we meta-analysed three separate age-at-sampling EWASs run within each OFC group (more details in Additional File 1), and the fact that many of these CpGs have previously been shown to be differentially methylated with age in infancy [31]. Of the 29,984 age-associated CpGs, 214 were also associated with CPO compared to CLO with P<1*10^− 7^. When we ‘filtered out’ the 29,984 age-related CpGs by setting the P-values to 1 in the CPOvsCLP and CPOvsCLO results, 121 CpGs were associated with CPO compared to CLO (P-value<l*10^−7^; Table 2; Additional File 2 Table S1). All subsequent analyses were performed on these results, that is, CLOvsCLP without filtering age-associated CpGs, and CPOvsCLP and CPOvsCLO with age-associated CpGs filtered. Full EWAS results for all three OFC subtype comparisons are available as additional files (Additional Files 3, 4 and 5) and summarised in Figure 3.

**Figure 3.**
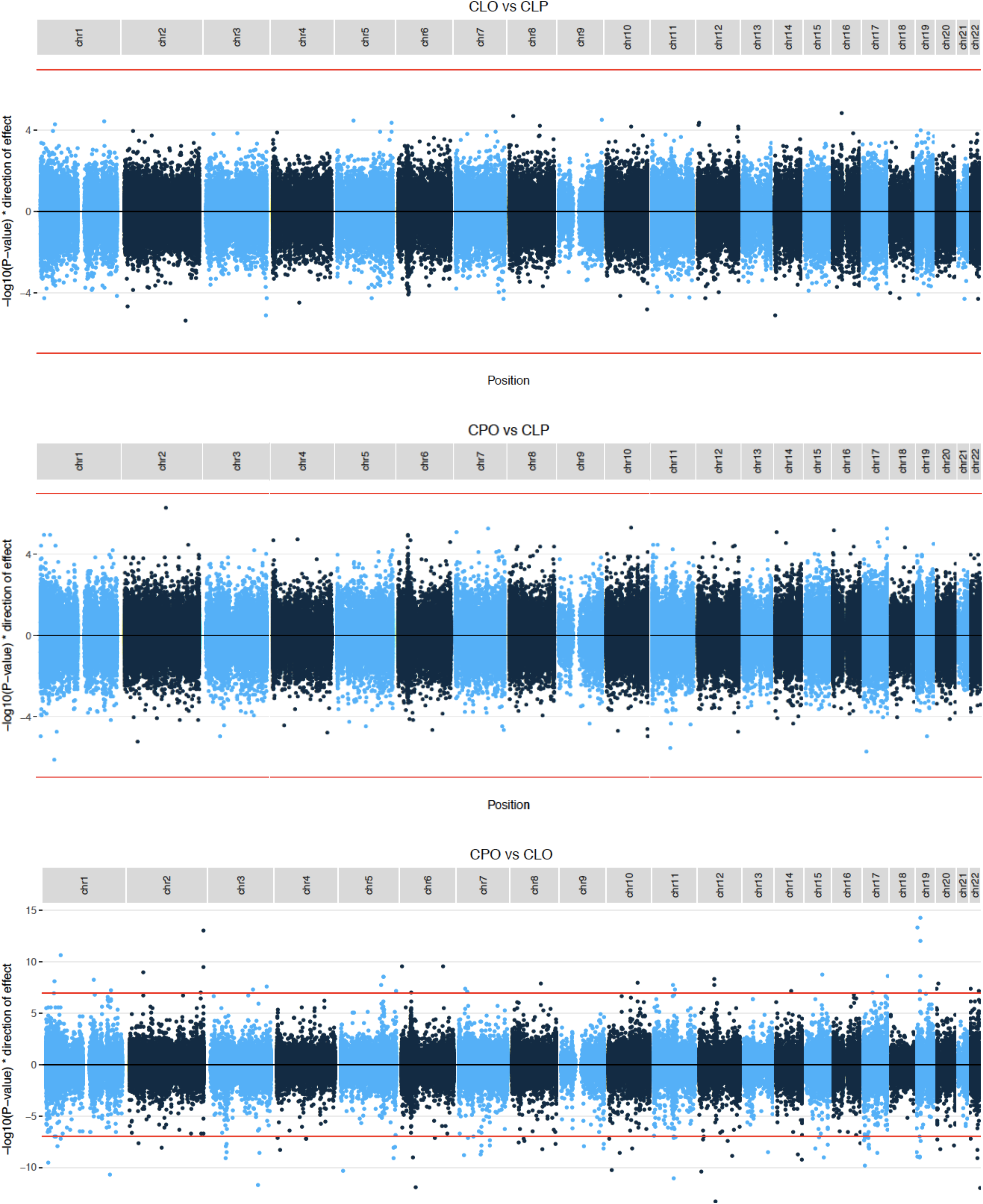
Manhattan plots of results from epigenome-wide association studies comparing OFC subtypes. Manhattan plots of the three pairwise epigenome-wide studies of DNA methylation in whole blood samples from children with CLO, CLP and CPO. P-values for age-related CpGs have been set to 1 (i.e. –1og10 P-value of 0) in the comparisons involving CPO.

**Table 2.** CpGs associated with CPO compared to CLO in the single site EWAS analysis.

| Chr | CpG | Regression coefficient | P-value | Gene | Relation to CpG island<sup>a</sup> | Relation to gene<sup>a</sup> |
| --- | --- | --- | --- | --- | --- | --- |
| 19 | cg01634146 | 0.19 | $9.80 \times 10^{-13}$ | *NFIX* | S_Shelf | Body |
| 22 | cg12899065 | 0.16 | $3.98 \times 10^{-8}$ | *GP1BB;SEPT5* | Island | TSS1500;<br>3'UTR |
| 6 | cg14623715 | 0.14 | $2.46 \times 10^{-10}$ | *PDE7B* | | Body |
| 10 | cg02017450 | -0.14 | $5.93 \times 10^{-11}$ | *intergenic [SFTA1P]* | | |
| 8 | cg04364695 | -0.13 | $4.27 \times 10^{-8}$ | *ZMAT4* | | Body |
| 7 | cg22114489 | -0.13 | $4.60 \times 10^{-8}$ | *intergenic [CUX2]* | | |
| 1 | cg12697139 | -0.13 | $2.23 \times 10^{-11}$ | *intergenic [MIR205HG]* | | |
| 2 | cg19075787 | -0.13 | $8.44 \times 10^{-9}$ | *intergenic [LOC284998]* | | |
| 11 | cg17696044 | -0.13 | $8.61 \times 10^{-8}$ | *SHANK2* | | Body |
| 20 | cg19592472 | -0.12 | $5.04 \times 10^{-8}$ | *OXT* | Island | 1stExon;<br>5'UTR |
| 4 | cg14348967 | -0.12 | $5.12 \times 10^{-9}$ | *intergenic [DQ599898]* | | |
| 6 | cg00257775 | -0.12 | $9.45 \times 10^{-10}$ | *ZFAND3* | | Body |
| 3 | cg25938530 | -0.11 | $2.04 \times 10^{-8}$ | *ITIH1* | | TSS200;B<br>ody |
| 11 | cg12155547 | -0.11 | $9.96 \times 10^{-12}$ | *intergenic [NEAT1]* | S_Shelf | |
| 1 | cg18147098 | 0.11 | $5.44 \times 10^{-8}$ | *intergenic [ATF3]* | S_Shore | |
| 8 | cg19496364 | 0.10 | $1.16 \times 10^{-8}$ | *intergenic [LINC00535]* | | |
| 7 | cg27508620 | 0.10 | $3.67 \times 10^{-8}$ | *intergenic [SP4]* | | |
| 6 | cg25426302 | 0.10 | $9.90 \times 10^{-8}$ | *PPT2;PRRT1* | N_Shore | TSS1500 |
| 10 | cg20327845 | -0.10 | $6.66 \times 10^{-8}$ | *PFKP* | | Body |
| 3 | cg05581878 | -0.10 | $2.50 \times 10^{-9}$ | *intergenic [AK097161]* | | |
| 10 | cg19220719 | -0.10 | $7.02 \times 10^{-9}$ | *intergenic [C10orf11]* | | |
| 22 | cg13251842 | 0.09 | $6.67 \times 10^{-8}$ | *MIRLET7A3;MIRLET7B* | | TSS200;<br>TSS1500 |
| 19 | cg27392771 | 0.09 | $5.25 \times 10^{-15}$ | *NFIX* | S_Shore | Body |
| 6 | cg23279756 | -0.09 | $7.14 \times 10^{-8}$ | *intergenic [ARMC2]* | | |
| 2 | cg07644939 | 0.09 | $3.12 \times 10^{-10}$ | *SNED1* | S_Shore | Body |

The top 25 CpGs with the largest effect sizes and P-values $< 1 \times 10^{-7}$ are shown. For intergenic regions, the closest annotated gene is shown in square brackets.
<sup>a</sup> S\_Shore = South shore; S\_Shelf, = South shelf; N\_Shore = North shore; TSS1500 = 1500
base pairs from a transcription start site; TSS200 = 200 base pairs from a transcription start
site; UTR = untranslated region

### Differentially methylated region analysis

When we interrogated differential methylation over larger regions of the genome, we found four DMRs in CLO compared to CLP, 17 in CPO compared to CLP and 294 in CPO compared to CLO (Sidak-corrected P-value<0.05; Table 3). The top DMRs with Sidak P-values<0.05 and the largest effect sizes are presented in Table 3. Boxplots of methylation levels averaged over the top DMRs for each subtype comparison are shown in Figure 4.

None of the 25 CpGs in the four CLOvsCLP DMRs overlapped with CpGs in DMRs from the other two comparisons. Of the 82 CpGs in the 17 CPOvsCLP DMRs, 39 (48%) were also in the list of 1,063 CpGs in the 294 CPOvsCLO DMRs (Figure 5). CPO was associated with higher methylation relative to CLP or CLO at 18 of these 39 CpGs and lower methylation relative to CLP or CLO at the remaining 21/39 CpGs.

**Figure 4.**
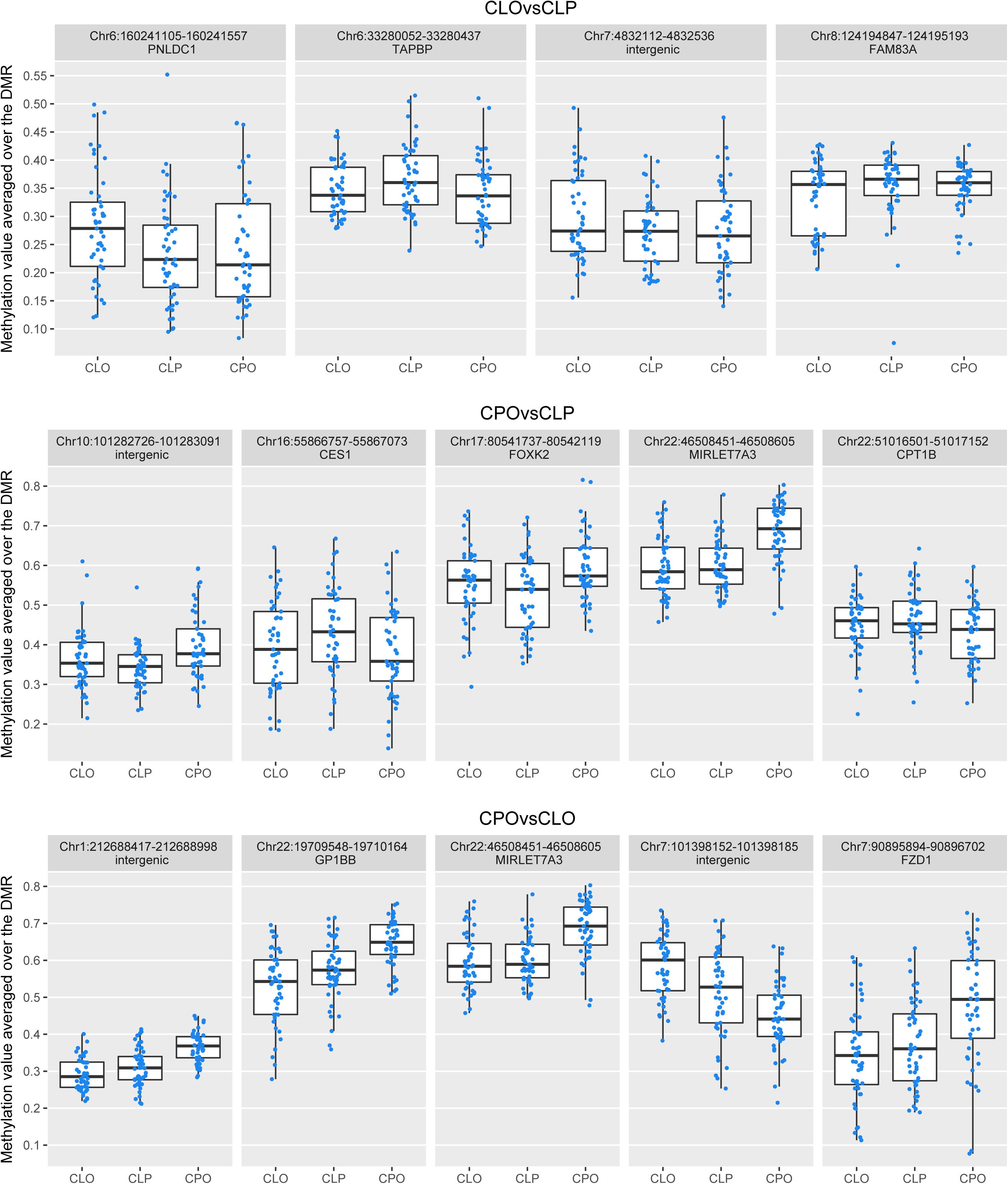
DNA methylation levels at the top differentially methylated regions. DNA methylation levels at the top DMRs (selected based on largest effect size and a Sidak-corrected P-value<0.05) for each pairwise epigenome wide study.

**Figure 5.**
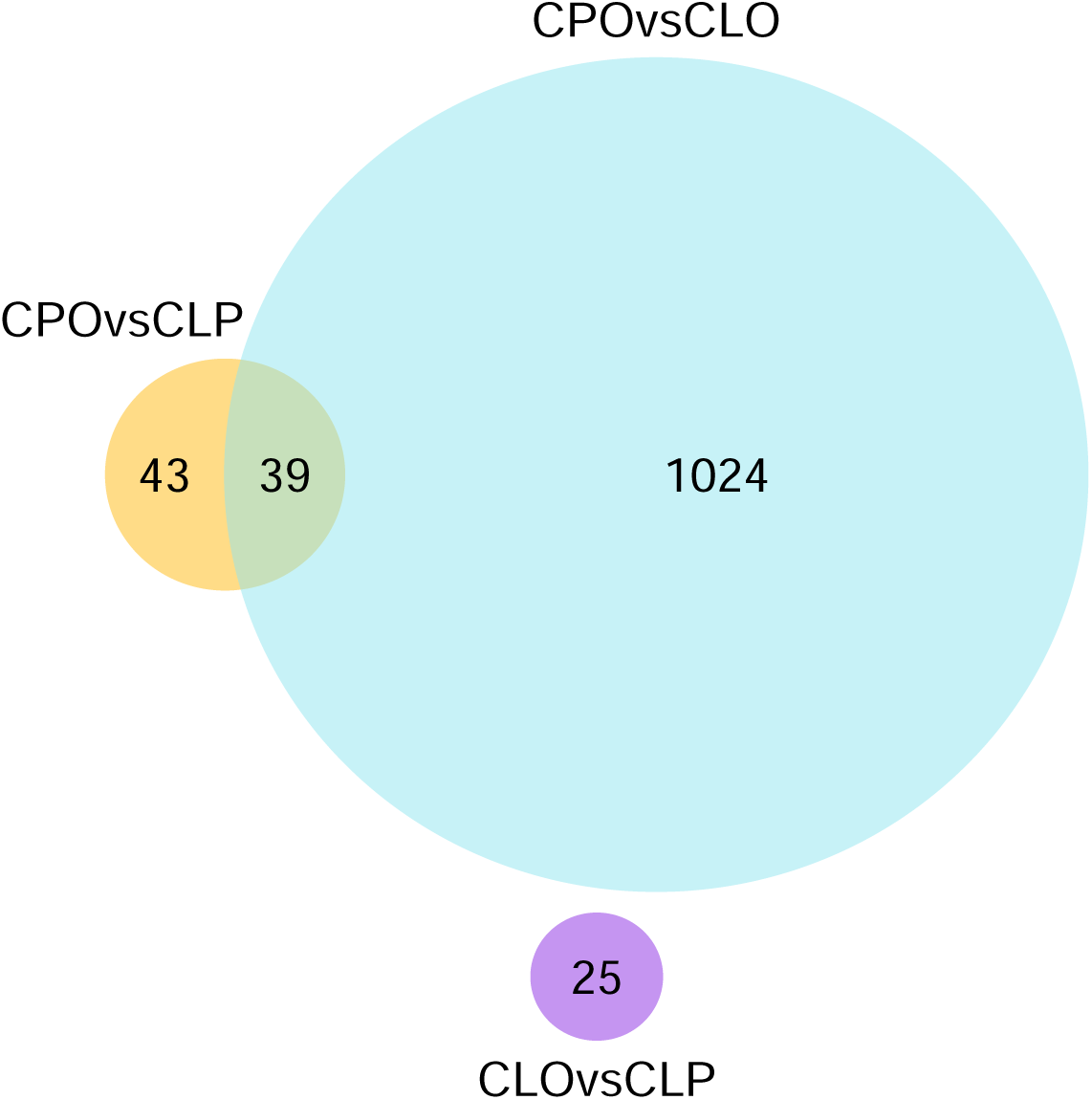
A Venn diagram to show the crossover in CpGs within DMRs associated with each subtype comparison.

**Table 3.** Top 5 DMRs with the largest effect sizes and Sidak-corrected P-values <0.05.

| **EWAS** | **DMR** | **Gene** | **N**<br>**CpGs** | **Sidak**<br>**corrected**<br>**P-value** | **Range of**<br>**regression**<br>**coefficients** |
| --- | --- | --- | --- | --- | --- |
| **CLOvsCLP** | Chr6:160241105-160241557 | *PNLDC1* | 4 | $4.8 \times 10^{-4}$ | 0.053, 0.078 |
| **CLOvsCLP** | Chr8:124194847-124195193 | *FAM83A* | 5 | $1.3 \times 10^{-3}$ | -0.072, -0.007 |
| **CLOvsCLP** | Chr6:33280052-33280437 | *TAPBP* | 11 | $2.8 \times 10^{-5}$ | -0.049, -0.008 |
| **CLOvsCLP** | Chr7:4832112-4832536 | *Intergenic*<br>[*KIAA0415*] | 5 | $8.6 \times 10^{-4}$ | 0.023, 0.059 |
| **CPOvsCLP** | Chr22:46508451-46508605 | *MIRLET7A3* | 6 | $8.07 \times 10^{-7}$ | 0.028, 0.119 |
| **CPOvsCLP** | Chr17:80541737-80542119 | *FOXK2* | 4 | $1.05 \times 10^{-6}$ | 0.060, 0.079 |
| **CPOvsCLP** | Chr10:101282726-101283091 | *Intergenic*<br>[*DQ372722*] | 5 | $7.23 \times 10^{-7}$ | 0.026, 0.077 |
| **CPOvsCLP** | Chr16:55866757-55867073 | *CES1* | 5 | $9.67 \times 10^{-5}$ | -0.093, -0.066 |
| **CPOvsCLP** | Chr22:51016501-51017152 | *CPT1B* | 12 | $1.47 \times 10^{-7}$ | -0.063, -0.026 |
| **CPOvsCLO** | Chr22:19709548- | *GP1BB* | 5 | $1.23 \times 10^{-14}$ | 0.067, 0.162 |
|  | 19710164 |  |  |  |  |
| **CPOvsCLO** | Chr22:46508451-46508605 | *MIRLET7A3* | 6 | $1.51 \times 10^{-14}$ | 0.035, 0.142 |
| **CPOvsCLO** | Chr7:101398152-101398185 | *Intergenic*<br>[*CUX1*] | 3 | $3.00 \times 10^{-13}$ | -0.131, -0.085 |
| **CPOvsCLO** | Chr1:212688417-212688998 | *Intergenic*<br>[*ATF3*] | 6 | $1.23 \times 10^{-16}$ | 0.018, 0.108 |
| **CPOvsCLO** | Chr7:90895894-90896702 | *FZD1* | 4 | $1.07 \times 10^{-9}$ | 0.114, 0.170 |

For intergenic regions, the closest annotated gene is shown in square brackets.

### Functional analysis of DMRs

There was no enrichment (FDR-adjusted P-value<0.05) for any functional categories defined using GO terms or KEGG pathways in CpGs within CLOvsCLP DMRs or CPOvsCLP DMRs. CpGs in CPOvsCLO DMRs were also not enriched for any GO terms, but were enriched for 66 KEGG pathways. However, these were mostly broad (for example, pathways in cancer, MicroRNAs in cancer, fatty acid metabolism). Results are presented in detail in the Additional File 2 Table S3.

According to the DisGeNET database, there were 286 genes associated with OFCs, of which 93 were also identified in a recent bioinformatics study [35] that found 357 unique OFC-related genes (total number of OFC-related genes from the literature: 643; Additional File S2 Table S4). Of these, eight mapped to regions that were differentially methylated in CPO compared to CLO in our study, and two mapped to regions that were differentially methylated in CPO compared to CLP. These genes are detailed in Table 4.

**Table 4.**
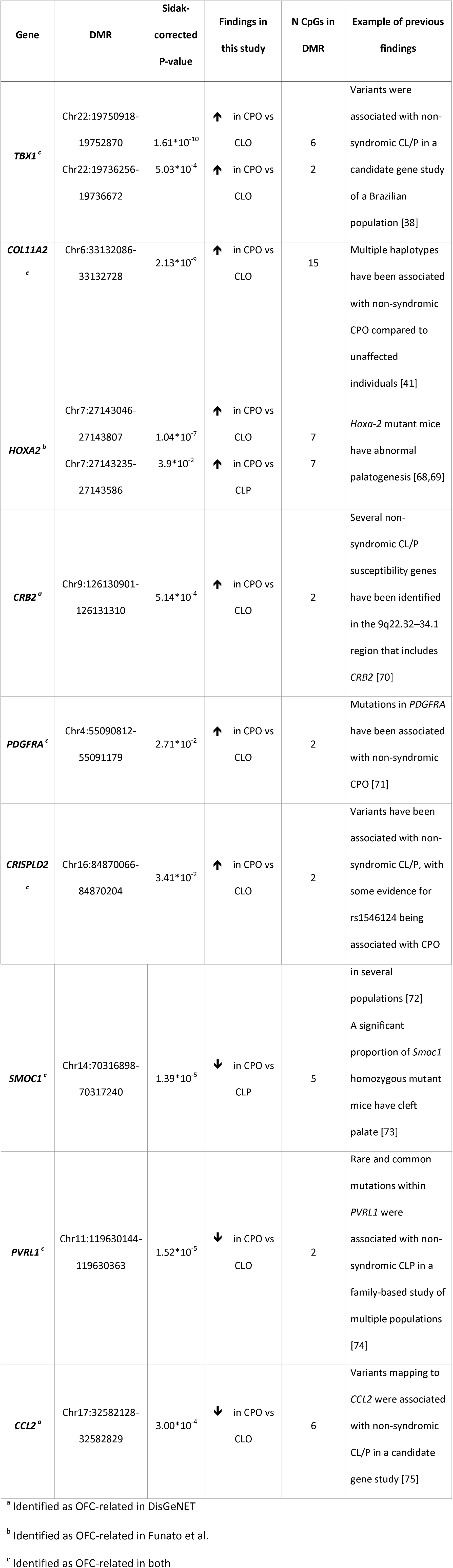
DMRs where genetic variation has previously been associated with OFCs.

## Discussion

In this, the first study of the epigenetic epidemiology of OFCs, we found multiple genomic regions differentially methylated in blood samples from non-syndromic children with CLO, CLP and CPO. Many more regions were differentially methylated between CPO and CLO than between CPO and CLP, and more regions were differentially methylated between CPO and CLP than between CLO and CLP. This suggests that the DNA methylation profiles of CLO and CLP are more similar to each other than the DNA methylation profile of CPO, which supports previous suggestions of distinct aetiologies for CPO and CL/P [5,36], Additionally, the DMRs we found between CLO and CLP support previous evidence that these subtypes are also aetiologically distinct. Our findings therefore have important implications for OFC research, suggesting that CLO, CLP and CPO should be analysed separately and not combined into a single entity or CL/P for analysis, at least in epigenetic studies.

There are three main possible explanations for why children with different OFC subtypes have different blood DNA methylation profiles:

Firstly, the subtypes might have distinct aetiologies in which DNA methylation plays a mechanistic role, i.e. the subtypes are caused, in part, by differences in DNA methylation. Arguably, for this to be the case, blood DNA methylation at the time of sampling (up to 20 months after birth) would have to closely reflect DNA methylation in the developing orofacial tissues during embryogenesis. For obvious ethical reasons, it is not possible to study these tissues in humans, however it is plausible that DNA methylation from postnatal blood might correlate with that from the developing embryo at some CpGs, particularly at metastable epi alleles where variable DNA methylation is established early in development before cellular differentiation.

Secondly, the subtypes might have distinct aetiologies explained by genetic and/or environmental factors that also influence blood DNA methylation. That is, the association between OFC subtype and DNA methylation is confounded by genotype or prenatal environmental factors such as maternal smoking or obesity. In this case, our finding of distinct blood DNA methylation profiles between subtypes would still support distinct aetiologies, but not a mechanistic role for blood DNA methylation.

Thirdly, since the OFC forms early in embryonic development and blood DNA methylation was measured in infancy, it is possible that the OFC subtype could indirectly influence blood DNA methylation, that is, any association between OFC subtype and DNA methylation could be explained by reverse causation. For (hypothetical) example, children with CPO or CLP might have more difficulty feeding compared to children with CLO and the subsequent different nutritional exposure may cause differences in DNA methylation between children with these subtypes. The Cleft Collective is currently collecting cord blood samples, which are unaffected by post-natal environmental factors, and will therefore help overcome this issue.

Further work is warranted to explore our findings in a causal analysis framework. However, several of our DMRs map to genes that have previously been associated with OFCs, which provides some support that DNA methylation either plays a causal role in development of OFC subtypes or reflects different genetic or environmental factors that do. For example, we identified six CpGs in a region on the gene body of *TBX1* that were 3 to 8% percent more highly methylated in blood DNA from children with CPO compared to CLO. *TBX1* encodes the T-box transcription factor 1 and deletion of this region causes chromosome 22q11.2 deletion syndrome, characterised by, amongst other malformations, cleft palate [37]. Genetic variants at *TBX1* have also been associated with non-syndromic CL/P [38]. *Tbx1* is expressed on the palatal shelves in mice and deletion results in abnormal epithelial fusion [39], We also identified a region of 15 CpGs on the gene body of *COL11A2* that were around 2% more highly methylated in CPO than CLO. *COL11A2* is one of three distinct genes that encodes collagen XI, which is expressed in the developing jaw in rats [40]. Genetic variants in *COL11A2* can cause syndromic and non-syndromic palatal defects [41,42].

Amongst our identified DMRs, there were some additional gene-OFC relations that have previously been reported in the literature, but that weren′t included in either the DisGeNET database or the recent review of OFC genes by Funato et al. [35]. For example, rare and/or common variants have been associated with non-syndromic CL/P at regions that were differentially methylated in CPOvsCLO: *FZD1* (hypermethylated) [43], *VAX2* (hypermethylated) [44] and *FGF12* (hypomethylated) [45]. We also found that children with CPO had lower methylation than children with CLO at a region of 5 CpGs near *MKNK2,* which has very recently been associated with non-syndromic CL/P in central Europeans [46], However this DMR did not survive correction for multiple testing (Sidak-corrected P=0.2).0ur finding of DMRs near OFC-implicated genes is consistent with the hypothesis that these loci play an important part in OFC aetiology, with two possible explanations for the observed associations: 1) they are explained by the underlying genetic architecture (and some of the children in our study may have undiagnosed syndromes caused by these genes); 2) they are explained by non-genetic variation in DN A methylation. Either way, these observations corroborate that perturbation of gene function at these loci is important in causing OFCs.

We also found over 250 novel genomic regions associated with different OFC subtypes, including four regions differentially methylated in CLO compared to CLP mapping to *PNLDC1, FAM83A, TAPBP* and an intergenic region with the nearest gene *KIAA0415.* Few genes have previously been implicated in CLO, because most studies have not considered it as molecularly distinct from CLP. Of the novel genes associated with CPOvsCLO or CPOvsCLP, we have selected a few that could be related to OFCs via a biologically plausible mechanism. For example, we found a region of six CpGs near *MIRLET7A3* that was 8% more highly methylated in CPO compared to CLO and 5% more highly methylated in CPO compared to CLP. *MIRLET7A3* encodes a microRNA precursor, and although the mechanistic role of microRNAs in human OFCs has not been fully explored, there is some evidence from mouse studies that they could be important [47]. Furthermore, a recent microarray study found that has-let-7a-5p, which is the mature sequence of the *MIRLET7A3-encoded* precursor, was over expressed in plasma samples from non-syndromic children with CPO and CLP relative to unaffected controls [48]. In our CPOvsCLO comparison, we also found several novel DMRs mapping to genes that have previously been linked to neural tube closure and/or defects (NTDs), for example *RGMA* [49], *ARHGEF1* [50] and *NODAL* [51], as well as two genes that have been linked to both OFCs and NTDs, *CCL2* [52] and *PDGFRA* [53]. This is particularly interesting, because NTDs and OFCs appear to share some aetiological features: They both occur when tissues in the midline fail to fuse completely during embryonic development [54]; they co-occur in the same individuals and in related individuals more than would be expected by chance [55]; they share several environmental risk factors [2,3,56,57]. Our findings further support recent evidence of an overlap in the molecular networks associated with OFCs and NTDs [58].

Finally, two of the DMRs we identified have previously been found in association with maternal risk factors for OFCs. A region of four CpGs at *HIF3A* was between 3 and 15% more highly methylated in children with CPO than in children with CLO. Methylation in this region has previously been associated with measures of adiposity, most commonly body mass index (BMI) [59], A previous study found a positive association between maternal BMI and offspring cord blood DNA methylation at the four CpGs in our *HIF3A* DMR [60]. Additionally, a region of two CpGs at *PRPH* was 5% more highly methylated in children with CPO than in children with CLO. Methylation at three (different) CpGs at *PRPH* has previously been negatively associated with maternal plasma folate levels [61]. These findings might indicate distinct aetiologies with different risk factors, suggesting that CPO and CLO are differentially influenced by maternal adiposity and/or maternal folate levels.

Although OFCs are one of the most common birth defects, they are relatively rare, so collecting data on large numbers of affected individuals is challenging [15]. We used data and samples collected as part of the Cleft Collective cohort study, which is a unique and valuable resource for OFC research. The prospective nature of this cohort means that future work can assess whether the subtype-associated methylation we see in infancy persists to later ages and is associated with longer term adverse outcomes of OFC such as poor educational attainment. Partly due to the novelty of these data and this resource, we were unable to find an independent cohort with similar data to replicate our findings. We hope to generate DNA methylation data for a larger sample of Cleft Collective children and test for replication in future studies. Another potential limitation of our study is that children with CPO were on average six to seven months older than children with CLO and CLP, which had a large influence on the results of the EWAS comparing these subtypes. Although we believe we were largely successful in our attempt to remove this influence by filtering out age-related CpGs using a very liberal P-value threshold of uncorrected P<0.05, there may be some residual influence. For example, three of the top 25 CpGs where there is most evidence of differential methylation between CPO and CLO (Table 2) map to two genes that have previously been reported as associated with gestational age and/or age in infancy: *NFIX* [30,31,62] and *SNED1* [31]. However, a previous microarray study of lip tissue found lower expression of *NFIX* in children with CLP compared to children with CLO even though both groups were sampled at 4-months-old, which provides some evidence that *NFIX* may be associated with OFCs independently of age [63]. Differences in the surgical protocol for lip and palate repair mean that this limitation (of age differences between children with CPO and CL/P) is likely to be present in other studies of OFCs where samples are collected at surgery, so techniques such as the one described in this paper should be developed to attempt to overcome this. We found no evidence of association between epigenetic age acceleration and OFC subtypes. Previous studies have postulated that epigenetic age acceleration is a measure of development in children [64,65], with a positive value indicating a child who is developmentally advanced for their actual age. Therefore our finding of no association suggests that children with different OFC subtypes have similar rates of development.

The Cleft Collective cohort is still in the recruitment stage and genotype and gene expression data do not yet exist for the participants. This means that we were not able to infer causality between OFC subtypes and blood DNA methylation using Mendelian randomization [66,67], or explore functionality by calculating correlations between methylation and expression. This is something we hope to do in further studies. There was a high proportion of missing demographic data (for example, on maternal smoking, education, occupation, ethnicity and parity), which is also related to the Cleft Collective cohort being in its infancy. Participants selected for this study were recruited near the start of the recruitment phase when questionnaire return rates were lower. Our return rates have increased recently and in future work, we hope to generate methylation data for a larger sample of the cohort with more complete questionnaire data.

The Cleft Collective is a case-only cohort, so we were unable to make comparisons with unaffected children. We explored several options for controls from other cohorts, but did not identify any options that would not have introduced significant confounding/bias by batch effects, age or tissue. When genotype data are available, future studies using Mendelian randomization will be able to circumvent these issues with confounding. Additionally, the Cleft Collective is collecting data on unaffected siblings, who will act as a good control sample in future studies.

In conclusion, we found several genomic regions differentially methylated in blood samples from non-syndromic children with CLO, CLP and CPO. Confidence in our results comes from the fact that many of these genes have been previously linked to OFCs, but we have also highlighted some novel regions. Our findings provide further support that CLO, CLP and CPO may be aetiologically distinct, which has important implications for future studies of OFC aetiologies.

## Declarations

### Ethics approval and consent to participate

Research ethical approval was granted by the NHS South West Central Bristol Ethics Committee (13/SW/0064). All participants provided informed consent to participate.

### Consent for publication

No individual level data is reported. Consent for publication of summary statistics was included in the research ethical approval (13/SW/0064).

### Availability of data and materials

To protect the privacy of participants in the Cleft Collective, we are unable to make individual level data available without a formal application for data access for specific projects. However, the datasets supporting the conclusions of this article (our full EWAS summary statistics) are available as additional files (Additional Files 3, 4 and 5).

## Authors’ contributions

GCS, CLR and GDS conceived and designed the study. AD, K.Humphries, SJL and JS manage recruitment and research within the Cleft Collective. K.Ho and WM processed the biological samples. AD selected participants from the Cleft Collective for this study. GCS normalised and quality controlled the methylation data. GCS analysed the data. ES and SJL provided advice on biological interpretation. All authors read and approved the final manuscript.

## Acknowledgements

This publication (project number CC002) involves data derived from independent research funded by The Scar Free Foundation (REC approval 13/SW/0064). We are grateful to the families who participated in the study, the UK NHS cleft teams, and The Cleft Collective team, who helped facilitate the study. We also thank the Bristol Bioresource Laboratory team for sample processing and storage.

